# Engineering Antibody-Drug Conjugates targeting an Adhesion GPCR, CD97

**DOI:** 10.1101/2025.08.24.671942

**Authors:** Takamitsu Hattori, Michelle Wang, Alexis D. Corrado, Suzanne Gross, Michelle Fang, Injin Bang, Nainita Roy, Iryna Berezniuk, Hayley Donaldson, Karenna Groff, Niklas Ravn-Boess, Akiko Koide, Dimitris G. Placantonakis, Christopher Y. Park, Shohei Koide

## Abstract

Adhesion G protein-coupled receptors (aGPCRs) are key cell-adhesion molecules involved in many cellular functions and contribute to human diseases, including cancer. aGPCRs are characterized by large extracellular regions that could serve as readily accessible antigens. However, the potential of aGPCRs as targets for biologic therapeutics has not been extensively explored. CD97, also known as ADGRE5, is an aGPCR that is upregulated in various cancer types, including acute myeloid leukemia (AML) and glioblastoma (GBM), and their respective cancer stem cells. Here, we developed antibody-drug conjugates (ADCs) targeting CD97 and assessed their efficacy against AML and GBM cells. We generated a panel of synthetic human antibodies targeting distinct epitopes of CD97, from which we identified an antibody that was efficiently internalized. This antibody binds to all isoforms of human CD97 but not to its close homolog, EMR2. Structure determination by single-particle cryo-electron microscopy revealed that this antibody targets the CD97 GPCR autoproteolysis-inducing (GAIN) domain, whose presence is conserved in aGPCRs, through an unconventional binding mode where it extensively utilizes the light chain framework for antigen recognition. Screening of conjugation methods and payloads resulted in a stable ADC that effectively killed AML and GBM cell lines, as well as patient-derived GBM stem cells, with minimal cytotoxicity against peripheral blood mononuclear cells from healthy donors. Our study demonstrates the therapeutic potential of targeting CD97, as well as the aGPCR GAIN domain in general, and broadens our mechanistic understanding of antibody-antigen interactions.

## INTRODUCTION

The adhesion G protein-coupled receptors (aGPCRs) comprise the second-largest class of the GPCR superfamily. An aGPCR consists of an N-terminal extracellular region (ECR), a seven-span transmembrane (TM7) region, and an intracellular region. It is characterized by the presence of a large ECR consisting of multiple structured domains, including diverse domains associated with protein-protein interaction and a characteristic GPCR autoproteolysis-inducing (GAIN) domain in proximity to the 7TM region (1-3). aGPCRs play key roles in physiology, including development and immunity, and are associated with human diseases such as neurodevelopmental disorders, immune disorders, and cancer (4-6). Unlike conventional GPCRs, which possess few well-structured regions on the extracellular surface and are primarily targeted with small-molecule drugs, the large extracellular regions of aGPCRs are attractive antigens for antibody-based therapies. However, their therapeutic potential remains largely unexplored (6). Indeed, there are currently no FDA-approved biologic drugs that target aGPCRs.

CD97, encoded by the *ADGRE5* gene, is a member of the epidermal growth factor 7-transmembrane (EGF-TM7) subfamily of aGPCRs (7, 8). Upregulation of CD97 occurs in immune cells during the inflammatory response, as well as in various types of malignant cells (9-16). CD97 is one of the most highly expressed antigens in AML cells, whereas its expression is lower in hematopoietic stem cells (HSCs) (11). It also plays a critical role in regulating blast growth, survival, and differentiation in AML. In glioblastoma (GBM), CD97 is expressed at much higher levels on tumor cells compared with normal brain tissue and neural progenitors, the putative cell-of-origin (12). CD97 is also overexpressed in leukemic stem cells (LSCs) and glioblastoma stem cells (GSCs) (11, 12). LSCs and GSCs contribute to therapy resistance and initiate relapse after treatment, and thus, their eradication is critical for improving treatment outcomes in AML and GBM (17, 18). Consequently, CD97 is a promising therapeutic target in AML and GBM.

CD97 has three distinct isoforms. Alternative splicing of CD97 mRNA leads to the inclusion or exclusion of different exons encoding EGF-like domains, resulting in the production of isoforms with varying numbers of EGF-like domains (**Fig.1**) (1, 8). Some of these EGF-like domains are responsible for ligand binding (1, 2). The GAIN domain is conserved across isoforms and is essential for the autocatalytic cleavage process as well as the receptor agonism caused by the exposure of the Stachel peptide segment (1-3). The remaining TM7 and intracellular regions play roles in interacting with G proteins and subsequent signal transduction. A closely related homolog, EGF-like module-containing mucin-like hormone receptor-like 2 (EMR2), shares essentially the same architecture as CD97, exhibiting high sequence identity in the EGF-like domains but moderate sequence identity in the GAIN domain with CD97 (**Fig.1**). Interestingly, despite their high sequence and structural similarity, CD97 expression is more frequently upregulated than EMR2 in AML and GBM cells, whereas EMR2 is mainly expressed in other hematopoietic cell types (11, 12, 19). Consequently, selectively targeting CD97 over EMR2 is preferable to reduce the risk of potential toxicity.

Many antibody-based therapeutic modalities harness immune cell-mediated cytotoxic mechanisms, including antibody-dependent cellular cytotoxicity (ADCC), antibody-dependent cellular phagocytosis (ADCP), and T cell-mediated responses elicited by immune checkpoint inhibitors, bispecific T-cell engagers, and chimeric antigen receptor (CAR) T-cells (20). As CD97 is upregulated in activated immune cells (9, 10), these strategies may not be able to effectively target CD97 expressed on the surface of cancer cells. Thus, we explored the utility of antibody-drug conjugates (ADCs) targeting CD97, as ADCs are a rapidly emerging class of therapeutic modalities that exert their effects independently of immune system engagement (21-23). ADCs deliver cytotoxic payloads selectively and directly into target cancer cells, improving the therapeutic index compared with conventional chemotherapies. ADCs consist of three main components: an antibody that targets a cell-surface antigen, a cytotoxic payload, and a cleavable or uncleavable linker that connects the antibody to the payload. A critical feature of the antibody component is that it binds to an antigen and promotes the internalization of the ADC-antigen complex into antigen-expressing cells. Thus, generating antibodies with high specificity and high internalization efficiency is crucial for developing potent ADCs.

Here, we describe the development of human synthetic antibodies against the extracellular region of human CD97. Among the antibodies we developed that target multiple distinct epitopes, we identified an antibody exhibiting high internalization efficiency suitable for generating ADCs. The cryo-EM structure revealed an unusual mechanism of this antibody that recognizes the CD97 GAIN domain. The ADC based on this antibody potently and selectively killed AML and GBM cells, demonstrating the therapeutic potential of targeting CD97 in these challenging cancers.

## RESULTS

### Generation and characterization of human synthetic antibodies targeting CD97

To generate antibodies that selectively recognize human CD97, we sorted a phage-display library of human synthetic antibodies using purified ECRs of CD97 isoforms 1 and 2, the longest and shortest of the isoforms (**Fig. 1A**). The selection process involved negative sorting using the ECRs of EMR2 isoforms (**Fig. 1A**). After library sorting, we produced candidate antibodies, A8, C27 and D25, in the Fab format and assessed their binding using a bead-based binding assay (24). All the antibodies exhibited apparent dissociation constant (*K*_D_) values in the nanomolar range for CD97 and did not bind to EMR2, indicating high affinity and selectivity (**Fig. 1B, Supplementary Fig. S1A**). While A8 and D25 bind to both CD97 isoforms, C27 binds to CD97 isoform 1 but not isoform 2 (**Fig. 1B, Supplementary Fig. S1A**). Epitope binning using competitive phage ELISA showed that their epitopes do not overlap (**Supplementary Fig. S1B**). We screened additional clones from this selection campaign, all of which were classified into one of the three already identified epitope bins.

**Figure 1.**
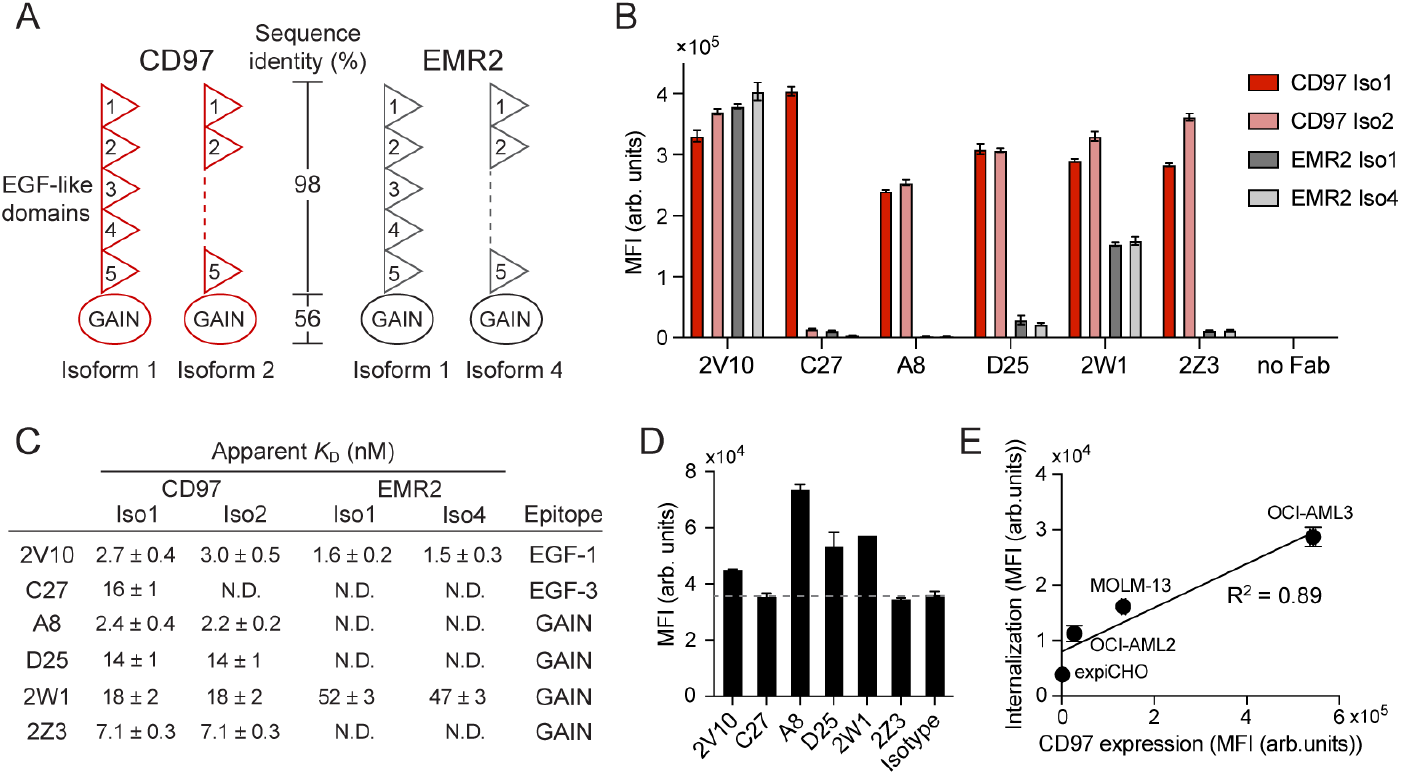
Development of human synthetic antibodies targeting CD97. (A) Schematic representation of the extracellular regions of CD97 and its close homolog, EMR2. (B) Binding analysis of anti-CD97 Fabs using a bead-based binding assay. The binding data obtained at the highest Fab concentration tested in the titration curves are shown. See Supplementary Fig. S1A for the binding titration curves. (C) The affinity and epitopes of anti-CD97 antibodies. The apparent dissociation constant (*K*_D_) values of the antibodies against each antigen were determined from the binding titration curves. The epitopes were determined by the epitope mapping assay using the truncation mutants of CD97. See Supplementary Fig. S1A and S2A for the original data. (D) Internalization of anti-CD97 antibodies into OCI-AML3 cells. The cells were incubated with the antibodies in the human IgG1 format with LALAPG mutations (IgG1-LALAPG) conjugated with the pHAb dye (500 nM) for 3 days. (E) The dependence of A8 internalization on the cell-surface CD97 level. The median fluorescence Intensity (MFI) obtained from the internalization assay was plotted against the MFI from the cell surface staining experiment using A8 for each cell type. Each MFI value was background-corrected by subtracting the corresponding value obtained from either the isotype control antibody-pHAb conjugate (for internalization) or the secondary antibody only control (for CD97 staining). For internalization assay, the cells were incubated with pHAb dye-conjugated antibodies (200 nM) for 24 hr. See Supplementary Fig. S2D for the original data. The mean and s.d. from N = 3 technical replicates are shown.

To maximize the likelihood of identifying antibodies with high internalization efficiency for developing effective ADCs, we generated additional antibodies that bind to epitopes other than the three described above. We conducted phage-display library sorting using OCI-AML3 cells, which express a high level of CD97, in the presence of saturating concentrations of A8, C27 and D25 Fab competitors to avoid recovering antibody clones that bind to the same epitopes as those antibodies. As expected, the epitopes of newly identified antibodies were distinct from the first three (**Supplementary Fig. S1C**). The second set of antibodies was classified into three epitope bins (**Supplementary Fig. S1D**). They also showed high affinity for CD97, but some of them exhibited different specificity profiles compared to the first set (**Fig. 1B, Supplementary Fig. S1A**). 2V10 bound to all CD97 and EMR2 isoforms tested, suggesting that it recognizes a region highly homologous between CD97 and EMR2. Together, we identified high-affinity antibodies that recognize a total of six non-overlapping epitopes on the CD97 ECR (**Supplementary Fig. S1E**).

To further define the epitopes of these antibodies, we designed truncation mutants of CD97 with systematic deletions of the EGF-like domains from the N-terminus, and assessed antibody binding to the cells expressing these variants (**Supplementary Fig. S2A**). The binding analysis revealed that 2V10 and C27 recognize the EGF-1 and EGF-3 domains, respectively, while the other antibodies recognize the GAIN domain. These data rationalize the selectivity of these antibodies among CD97 isoforms and EMR2 (**Fig. 1B,C, Supplementary Fig. S1A**). The EGF-1 domain is present across all CD97 isoforms and shares 95% sequence identity with that of EMR2, which explains the binding of 2V10 to all tested isoforms of both CD97 and EMR2 (**Fig.1B, Supplementary Fig. S1A**). The EGF-3 domain is absent in CD97 isoform 2, explaining selectivity of C27 for CD97 isoform 1 (**Fig.1A,B, Supplementary Fig. S1A**). Notably, C27 does not bind to EMR2, despite the EGF-3 domain sharing 93% sequence identity between CD97 and EMR2. These data demonstrate the feasibility of developing antibodies selective for each isoform of CD97. The sequence identity of the GAIN domain between CD97 and EMR2 is low (56%), rationalizing the selectivity of GAIN-targeting antibodies towards CD97 and our success in finding more GAIN binders than EGF binders from our antibody discovery campaign that included negative selection using EMR2.

To assess the affinity and internalization efficiency of antibodies using a format commonly used for ADCs, we produced six antibodies in the human IgG1 format. To eliminate target-independent binding to cells, we introduced the L234A/L235A/P329G (LALAPG) mutations into the Fc region (25), abolishing interaction with Fc receptors expressed on some immune cells as well as AML cells (26). Indeed, the isotype control antibody in the human IgG1 format strongly bound to OCI-AML3 cells, whereas the same antibody with LALAPG mutations showed negligible binding to these cells, which express Fc receptors (**Supplementary Fig. S2B**). Unless otherwise stated, the subsequent studies were performed using antibodies in the human IgG1-LALAPG format. Binding analysis using OCI-AML3 cells revealed that all antibodies exhibited EC_50_ values in the nanomolar range (we use EC_50_ instead of *K*_D_ because the binding measured here was likely mediated by bivalent interactions), indicating high affinity for endogenous CD97 expressed on the cell surface (**Supplementary Fig. S2C**).

We then evaluated the internalization efficiency of the antibodies. We conjugated the antibodies with the pHAb dye, which is pH-sensitive and exhibits low or no fluorescence at a pH greater than 7, but becomes fluorescent upon internalization and acidification in endosomes and lysosomes. Incubation of OCI-AML3 cells with the conjugated antibodies at saturating concentrations showed that they exhibited varying degrees of fluorescence enhancement, even among those targeting the GAIN domain, indicating that distinct epitopes within the same domain are associated with differences in internalization efficiency (**Fig. 1D**). A8 exhibited the highest internalization among the antibodies tested. Its internalization efficiency generally correlated with the expression level of CD97 when tested against a panel of cell lines (**Fig. 1E**), consistent with internalization of A8 being mediated by antigen binding. Together, we have identified an antibody targeting CD97 well-suited for ADC application.

### Cryo-EM structure of A8 in complex with CD97

To elucidate the molecular basis for the recognition of CD97 by A8, we determined the cryo-EM structure of A8 Fab in complex with the ECR of CD97 isoform 1 at a nominal resolution of 3.1Å (**Fig.2A, Supplementary Fig. 3**,**4**). We introduced the S531A mutation in the GAIN domain to prevent autoproteolytic cleavage at the G-protein coupled receptor proteolytic site (GPS) (27). We observed a clear map for the A8 variable domains and the CD97 GAIN domain, but not for the EGF-like domains, likely due to the high flexibility of the EGF-like domains (**Supplementary Fig. 3**,**4**). Our CD97 GAIN structure (**Fig. 2A**) is similar to the previously reported cryo-EM structure (PDB ID:8IKJ, RMSD for Cα = 0.874 Å; **Supplementary Fig. 5A)**, as predicted.

**Figure 2.**
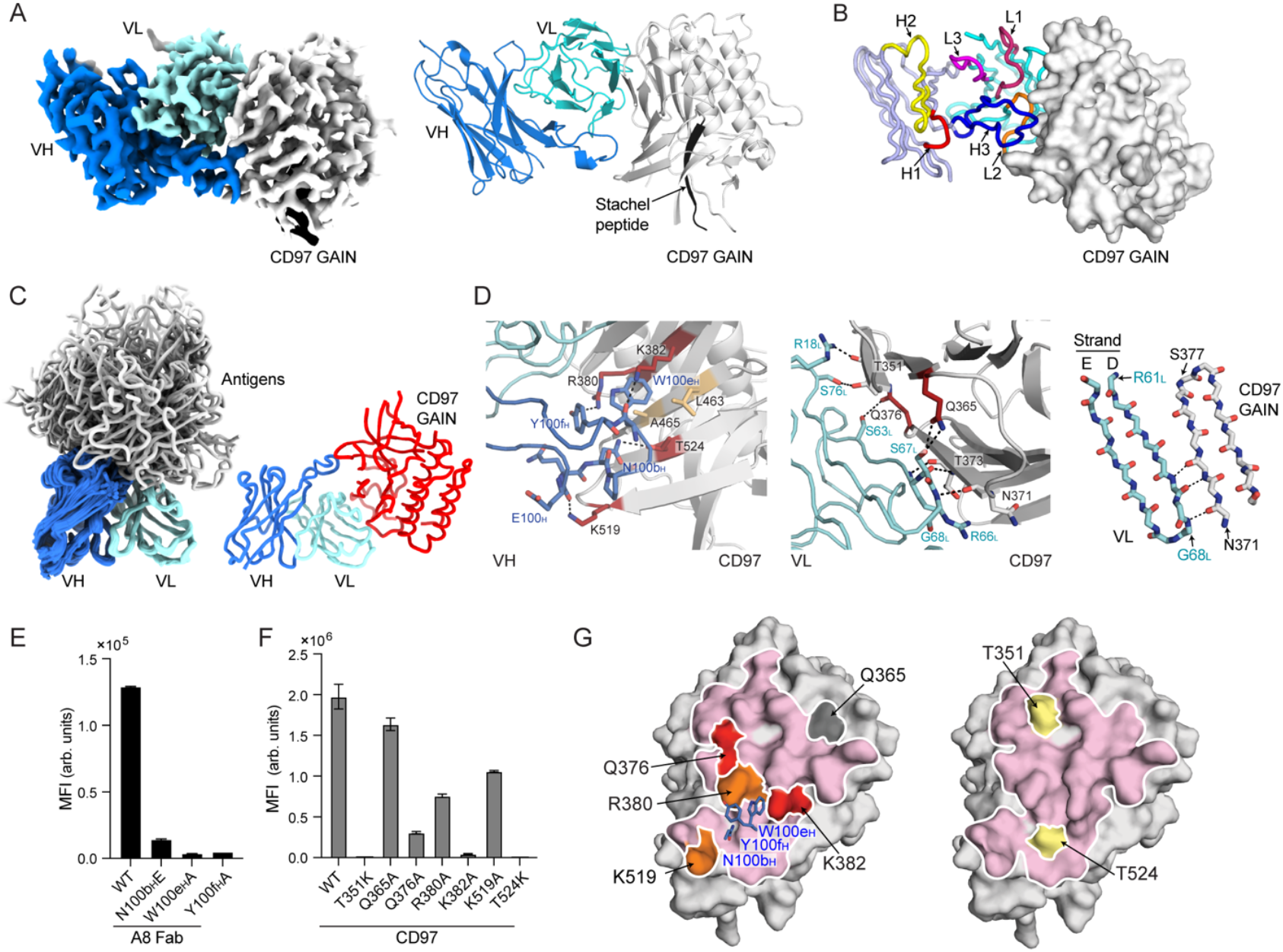
Cryo-EM structure of A8 Fab in complex with CD97 and mutation studies of interface residues. (A) Cryo-EM structure of A8 bound to CD97 isoform 1. Only the variable domains of A8 and the GAIN domain of CD97 are shown for clarity. The density map (left panel) and the structure (right panel) are shown. (B) The locations of the A8 CDRs in the A8-CD97 complex. (C) Comparison of the A8-CD97 complex to other antibody-antigen complexes. The left panel shows an overlay of a total of 43 published structures fulfilling the following criteria: a resolution below 2.5 Å, the use of the VH3 and Vk1 framework as in A8, and a human globular protein as the antigen. Portions of the antigens that do not directly interact with the antibodies have been omitted for clarity. (D) Close-up views of the interfaces between A8 VH and CD97 (left) and between A8 VL and CD97 (center). The residues involving interaction are labeled and shown in the stick model. The backbone atoms for a portion of the intermolecular β-sheet are shown in the right panel. Polar interactions are marked as dashed lines. (E and F) Binding of A8 mutant Fabs (6 nM) to CD97 isoform 1 immobilized on beads (E) and A8 Fab at a concentration of 40 nM to CD97 GAIN mutants expressed on the expiCHO cells (F) measured using flow cytometry. The mean and s.d. from N = 3 technical replicates are shown. See Supplementary Fig. S5B and S5C for the complete datasets with the titration curves. (G) The locations of CD97 residues tested using mutagenesis. The A8 binding interface is outlined on the CD97 surface, and the residues mutated to Ala (left) and to Lys (right) are indicated. Surfaces in red, orange and yellow indicate residues exhibiting significant reduction in binding upon mutation. The side chains of key residues in CDR H3 in A8 are also displayed as the stick model (left).

**Figure 3.**
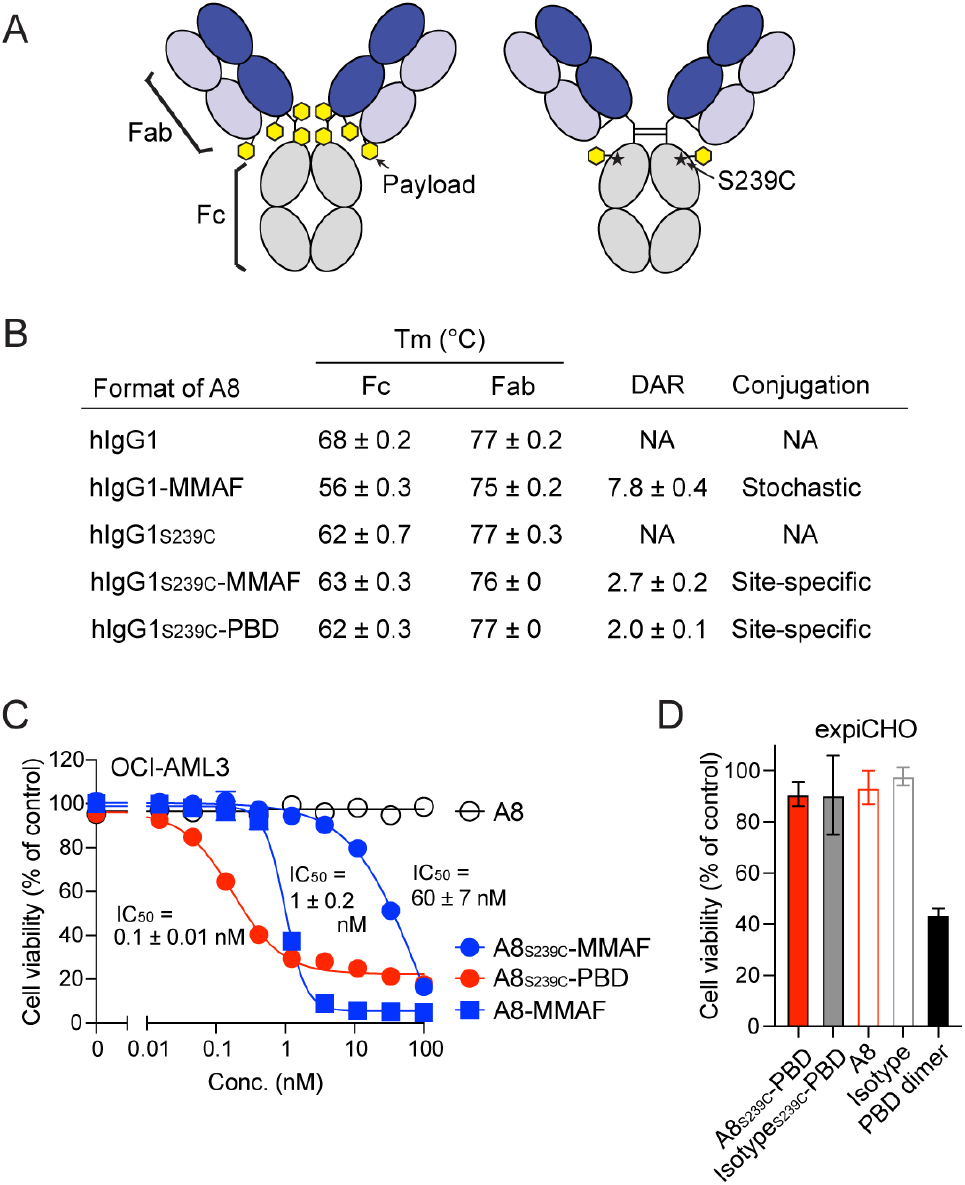
Development of ADCs and analysis of their cytotoxic effect. (A) Schematic representation of A8 ADCs prepared with the cysteine-based random (left) and site-specific (right) conjugation methods. Note that both formats contain the LALAPG mutations. (B) Thermal stability, drug-to-antibody ratio (DAR) and conjugation method of ADCs. See Supplementary Fig. S6 for original data. (C) Dose-response curves of ADCs using OCI-AML3 cells. The cells were incubated with the indicated ADCs or unconjugated A8 for four days, and cell viability was measured. The IC_50_ values are shown. (D) Cytotoxicity of A8_S293C_-PBD using expiCHO cells that do not express human CD97. The cells were incubated with either 100 nM ADCs, 100 nM unconjugated antibodies, or 10 nM PBD dimer for four days. The live cell percentages relative to the PBS control are shown. The mean and s.d. from N = 3 technical replicates are shown.

The cryo-EM structure revealed that A8 binds to the CD97 GAIN domain in an unconventional manner. Unlike conventional antigen-binding surfaces formed primarily by the complementarity-determining regions (CDRs), the binding interface of A8 is mainly created by the VL framework region along with portions of CDRH3 and L2 (**Fig. 2A, B**). The other four CDRs do not make direct contacts with the antigen (**Fig. 2B**). A structural comparison with other antibody-antigen complexes of antibodies with the same frameworks as A8 (VH3 and Vκ1) highlights the unique binding mode of A8, where the CD97 GAIN domain is located “on the side” of the VL domain, away from the surface produced by the CDRs to which most antigens bind (**Fig. 2C**). Interestingly, the exposed edge of the D strand of VL interacts with CD97 GAIN to form an intermolecular β-sheet (**Fig. 2D**). R18_L_ (subscript L denotes residues in the VL domain) in framework 1 (FR1) and S63_L_, R66_L_, S67_L_, G68_L_, and S76_L_ in framework 3 (FR3) of VL are involved in the formation of this intermolecular β-sheet. Additionally, the long CDRH3 extends over the VL domain and interacts with the β-sheet surface of CD97 GAIN primarily using E100_H_, N100b_H_, W100e_H_ and Y100f_H_ (**Fig. 2D**). We will further discuss this unusual binding mode below in Discussion.

We validated the cryo-EM structure by examining the effect of specific mutations in A8 predicted to disrupt its interactions with CD97 (**Fig. 2E-G, Supplementary Fig. S5B**,**C**). The mutations at N100b_H_, W100e_H,_ and Y100f_H_ in A8 CDRH3 reduced binding to CD97, indicating that these residues are essential for the A8-CD97 interaction (**Fig.2E,G, Supplementary Fig. S5B**). Similarly, K382A and T524K in the CD97 GAIN domain reduced binding to A8, confirming the structure (**Fig. 2F,G, Supplementary Fig. S5C**). Furthermore, the Q376A and T351K mutations within the CD97 GAIN domain intended to disrupt the interaction with the VL framework, diminished binding to A8, highlighting a critical role of the intermolecular β-sheet in A8 binding (**Fig. 2F,G, Supplementary Fig. S5C**). Notably, critical residues within CD97 involved in A8 interaction are not conserved in EMR2, explaining the lack of binding of A8 to EMR2 (**Supplementary Fig.S5D**). Collectively, the cryo-EM structure revealed the molecular basis for the recognition of the CD97 GAIN domain by the A8 antibody.

### Development of anti-CD97 ADCs

To develop ADCs based on the A8 antibody, we conjugated monomethyl auristatin F (MMAF), a tubulin inhibitor commonly used for ADCs (28), with a cleavable linker to A8 IgG using a cysteine-based random conjugation method (termed A8-MMAF; **Fig. 3A, Supplementary Fig. S6A**). A8-MMAF had a drug-to-antibody ratio (DAR) of 7.8 as estimated using MALDI-TOF mass spectrometry analysis (**Fig. 3B, Supplementary Fig. S6B)**, suggesting that all available cysteine residues after the reducing inter-chain disulfide bonds (a total of 8) were conjugated with MMAF. It showed cytotoxicity against OCI-AML3 cells with an IC_50_ of 1.0 nM (**Fig. 3C**), and selectively killed GBM cells that express CD97 (12). Although A8-MMAF exhibits effective cell-killing, its thermal stability was much lower than that of the unconjugated A8 sample (**Fig. 3B, Supplementary Fig. S6C**). SDS-PAGE under non-reducing conditions showed two major bands, consistent with the unpairing of disulfide bonds between the chains of A8-MMAF, suggesting that the reduction of inter-chain disulfide bonds contributed to this reduced stability (**Supplementary Fig. S6D**). The disruption of inter-chain disulfide bonds had a more pronounced effect on the stability of the Fc region than the Fab region (**Fig. 3B, Supplementary Fig. S6C)**, consistent with a previous study that reported the thermal stability of ADCs decreased as the average DAR increased, particularly in the CH2 domain (29).

To improve ADC stability, we introduced the previously described S239C mutation into the CH2 domain in Fc for site-specific conjugation (**Fig. 3A**) (30). This approach enables the production of homogeneous ADC with a narrow range of DAR while preserving the inter-chain disulfide bonds. As expected, an ADC sample produced using this method, A8_S239C_-MMAF, exhibited thermal stability comparable to that of unconjugated A8_S239C_ with a DAR of approximately 2 (**Fig. 3B, Supplementary Fig. S6A-D**). However, the cell-killing efficacy of A8_S239C_-MMAF was lower than A8-MMAF, likely due to the lower DAR of A8_S239C_-MMAF (**Fig. 3C**). To compensate for the low DAR, we utilized pyrrolobenzodiazepine (PBD) dimer, a potent DNA-crosslinking drug, as a payload. PBD dimer has a potential advantage in targeting cancer stem cells (CSCs) over mitotic inhibitors such as MMAE and MMAF, because CSCs can enter a quiescent state in which they cease proliferating, making mitotic inhibitors less effective (31, 32). Notably, this ADC, termed A8_S239C_-PBD, showed higher efficacy than both versions of the MMAF conjugates (**Fig. 3C**). A8_S239C_-PBD did not kill expiCHO cells lacking human CD97 (**Fig. 3D**), consistent with the notion that the cytotoxicity of CD97-ADC depends on CD97 expression. Taken together, we have generated a highly stable and potent anti-CD97 ADC.

### Cytotoxicity of anti-CD97 ADC against AML and GBM cells

Next, we evaluated the efficacy of A8_S239C_-PBD against a panel of AML cell lines. Staining of cells with A8 revealed that the CD97 level varied among these AML cells, with OCI-AML3 and HL60 showing the highest levels (**Supplementary Fig. S7A)**. A8_S239C_-PBD showed potent cytotoxicity against two CD97-high cell lines, OCI-AML3 and NB4, with IC_50_ values of 0.06 nM and 0.07 nM, respectively (**Fig. 4A**). In contrast, the IC_50_ for HL60 cells was much higher than the other AML cell lines, despite their high CD97 levels (**Fig. 4A)**. HL60 cells were much less sensitive to PBD dimer than other AML cells (**Supplementary Fig. S7B**), consistent with their known multidrug resistance (33). A8_S239C_-PBD exhibited approximately 8 to 30 times weaker killing efficacy against MOLM-13, KG-1a, and OCI-AML2 compared with OCI-AML3, likely due to the combination of their lower surface CD97 levels and differences in payload sensitivity (**Fig. 4A, Supplementary Fig. S7B**). The cytotoxicity of A8_S239C_-PBD was greater than that of Isotype_S239C_-PBD for all CD97-positive cells tested, indicating that the cell killing by A8_S239C_-PBD depends on CD97 expression (**Fig. 4A)**. The cytotoxic effect observed with Isotype_S239C_-PBD is likely due to antigen-independent cellular uptake, such as pinocytosis, or to the release of the payload outside the cells, which commonly occurs in ADCs (23).

**Figure 4.**
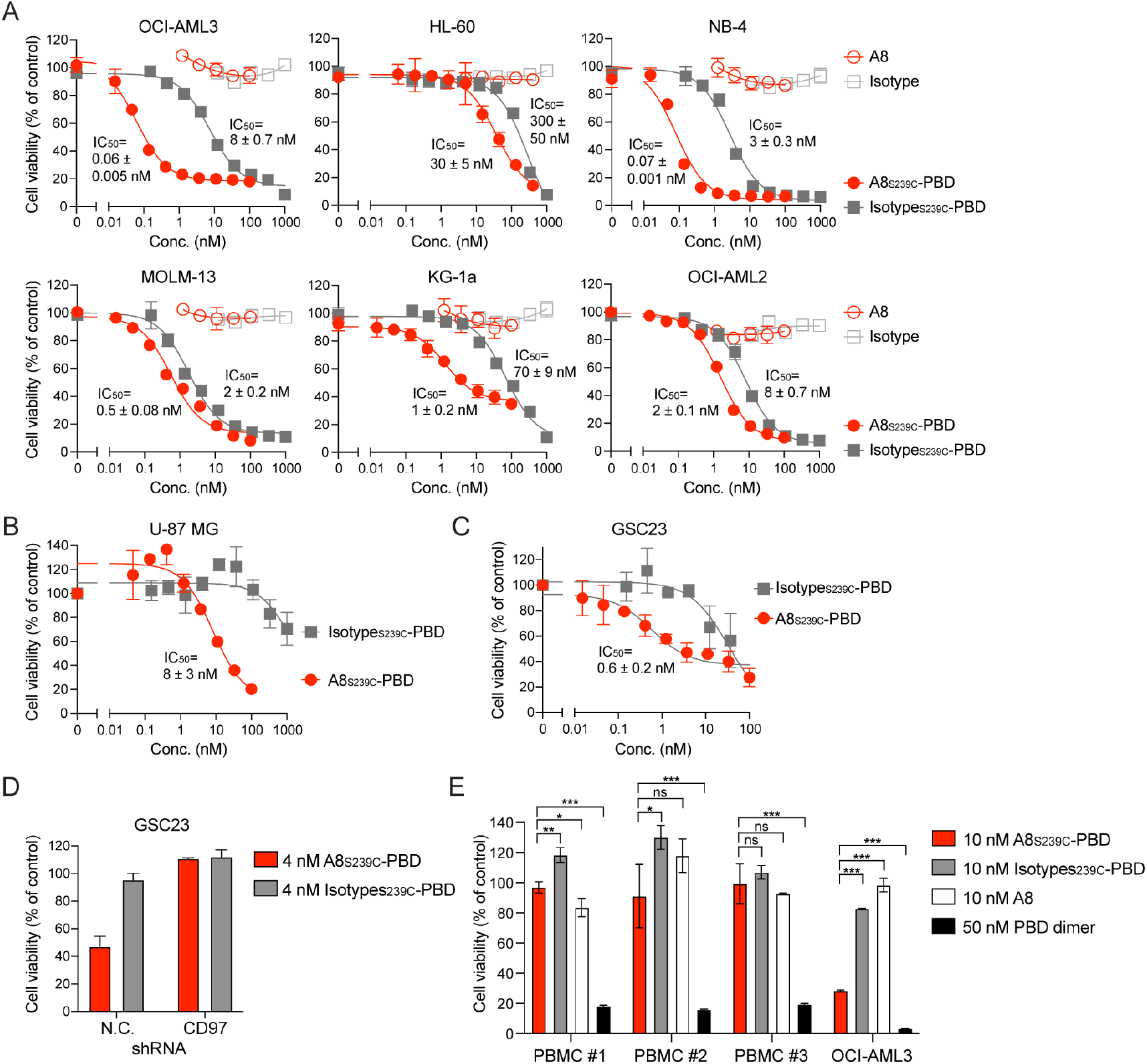
Analysis of the cytotoxicity of anti-CD97 ADC. (A-C) Dose-response curves of A8_S239C_-PBD and controls against a panel of AML cell lines (A), U-87 MG GBM cells (B) and GSC23 patient-derived GBM stem cells (C). The calculated IC_50_ values are shown. See Supplementary Fig. S7 and S8 for the CD97 levels of these cells. (D) Effect of A8_S239C_-PBD on GSC23 cells with and without CD97 knockdown using shRNA. (E) Toxicity of A8_S239C_-PBD on peripheral blood mononuclear cells (PBMCs). PBMCs from three healthy donors were incubated with the indicated reagents for four days, and cell viability was measured. OCI-AML3 cells were used as a positive control for A8_S239C_-PBD. The live cell percentages relative to the PBS control are shown. See Supplementary Fig. S9 for the complementary experiment measuring apoptosis in PBMCs. Data represent mean ± s.d. (N = 3), one-way ANOVA with Tukey’s multiple comparison test; *P < 0.05, **P < 0.01, ***P <0.001.

We further evaluated the efficacy of A8_S239C_-PBD against GBM cells. It showed greater cytotoxicity against U-87 MG cells than Isotype_S239C_-PBD, confirming CD97-mediated cytotoxicity (**Fig. 4B, Supplementary Fig. S8A**). Moreover, it showed potent cytotoxicity against patient-derived GBM stem cells, GSC23, which expressed high levels of CD97 on the surface (**Fig. 4C, Supplementary Fig. S8A-C**). The cytotoxicity of A8_S239C_-PBD was abolished upon CD97 knockdown using shRNA (**Fig. 4D, Supplementary Fig. S8B**), validating the antigen-dependent effects of the ADC. These data demonstrate the selective cytotoxicity of A8_S239C_-PBD against CD97-positive cells and its ability to effectively target patient-derived GBM stem cells.

Finally, we assessed the toxicity of A8_S239C_-PBD against healthy donor peripheral blood mononuclear cells (PBMCs) that include lymphocytes, monocytes, and dendritic cells. Treatment with A8_S239C_-PBD at 10 nM, a concentration ∼170 times higher than its IC_50_ against OCI-AML3 cells, induced only small or insignificant decreases in PBMC cell viability (**Fig. 4E, Supplementary Fig. S9**), despite the fact that monocytes, lymphocytes, and dendritic cells express low levels of CD97 (9, 34). These data indicate that A8_S239C_-PBD exhibits low toxicity to endogenous immune cells and suggest that A8_S239C_-PBD would exhibit low toxicity if administered to patients.

## DISCUSSION

In this study, we developed ADCs targeting CD97, an aGPCR, which belongs to a class in the GPCR superfamily whose therapeutic potential remains underdeveloped. Previous studies demonstrated the feasibility and efficacy of antibodies targeting aGPCRs including EMR1 for eosinophilic disorders (35), CD97 for arthritis (36), and ADGRG2 for testicular inflammation (37). In the context of cancer, GPR56 has been explored as a therpauetic target in colorectal cancer (38), and we and others have demonstrated the therapeutic potential of targeting CD97 in GBM (12, 39). This study presents a new avenue for targeting CD97 in AML and validates its therapeutic potential in GBM, further exemplifying and highlighting the potential of targeting aGPCRs using antibody-based strategies.

Targeting cancer stem cells such as LSCs and GSCs is crucial for achieving improved treatment outcomes, as they are key drivers of therapeutic resistance and relapse (17, 18). We showed effective cytotoxicity of our anti-CD97 ADC against patient-derived GBM stem cells. Previous studies reported the development of ADCs toward either LSCs or GSCs, including ADCs targeting CD33, CLL-1, CD123 and FTL3 for LSCs (32, 40-42), and EVA-1 for GSCs (43). Unlike these antigens, CD97 is highly expressed on both LSCs and GSCs (11, 12), allowing CD97-directed ADCs targeting both AML and GBM, two distinct types of cancer that depend on CD97 for stem cell maintenance. CD97 is also highly upregulated in other types of malignancies, including colorectal, thyroid, gastric and prostate carcinomas (13-16). Thus, we anticipate that CD97-directed ADCs could be utilized to treat a diversity of cancers.

The discovery of effective targeting of the GAIN domain of CD97 has important implications for drug discovery against other aGPCRs. The ECRs of aGPCRs have diverse compositions consisting of different modular domain families, but the GAIN domain is the only domain conserved among all aGPCRs (1, 4). We speculate that antibodies targeting the GAIN domain may facilitate the antigen-mediated internalization of other aGPCRs and that such antibodies may serve as effective starting materials for developing ADCs. Future studies will test this hypothesis.

While the focus of this study is ADC development using a single antibody, other antibodies generated in this study are potential therapeutic candidates and/or research tools for studying CD97 function. For example, they can be used to construct biparatopic antibodies, designed to bind two distinct epitopes of a protein, that can exhibit greater internalization efficacy than standard antibodies that target a single epitope (44). Additionally, previous studies have shown functional modulation of aGPCRs using antibodies and synthetic binders, which trigger allosteric conformational changes (37, 45) or inhibit ligand binding (46). A series of antibodies developed in this study potentially modulate CD97 functions through similar mechanisms. Future studies will explore these possibilities.

The structure of the A8-CD97 ECD complex (Fig. 2) uncovered an unconventional mode of antibody-antigen interaction. Conventionally, paratope residues, the residues that interact with the antigen, are predominantly located in the CDRs of both heavy and light chains. Indeed, Madsen et al. analyzed 1,058 non-redundant structures of antibody-antigen complexes and revealed that over 80% of paratope residues are found in CDRs (47). In contrast, the paratope residues of A8 are mainly located in framework 3 of the light chain (LFR3) and CDRH3, with a few additional residues in LFR1 and CDR L2, resulting in only 50% of the paratope residues of A8 being in CDRs. Among the 1,058 antibodies in the studies by Madsen et al., residues equivalent to the paratope residues in LFR1 and LFR3 of A8 are rarely involved in antigen binding (less than 2% at R18_L_, S63_L_, S67_L_, G68_L_ and S76_L_) (47). Despite this unconventional binding mode, the binding interface area between A8 and CD97 is 1,063 Å^2^ (**Supplementary Table 1**), similar to the common binding interface area found in antibodies (48). It is intriguing that paratope residues in LFR1 and LFR3 of A8 are derived from the residues preserved in the germline (IGKV1-39*01), except for R66_L_, which is a rare amino acid residue at this position and is inherited from Trastuzumab used as the scaffold in our synthetic antibodies. The isotype control antibody harboring the same residues at these positions does not bind to CD97, suggesting that the interaction between A8 VL and CD97 GAIN is weak, but the cumulative effect derived from CDR H3 significantly enhances the overall affinity between A8 and CD97 GAIN.

The formation of an intermolecular β-sheet between the D-strand in LFR4 of A8 and the β-strand in the CD97 GAIN domain (from N371 to S377) is uncommon in antibody-antigen interactions (**Fig. 2D**), as paratopes are typically composed of loop structures formed by CDRs. The intermolecular β-sheet formation between antibodies and antigens has been observed in anti-HIV-1 gp120 antibodies (49, 50). However, in these antibodies, the β-strands formed within the extended CDR H3 interact with the β-strands in the antigens, clearly distinct from the binding mode observed in the A8-CD97 complex. These examples further highlight the unconventional antigen recognition by A8. We anticipate that these insights could be potentially valuable for designing new types of antibodies in which the framework region contributes to antigen binding.

## MATERIALS AND METHODS

Antibody selection using phage display was performed as described previously (51). Antibody production and characterization, including bead-based and cell-based binding assays, were performed according to published methods (52). Cryo-EM sample preparation, data collection, and processing were conducted as described previously (53). ADC production and characterization, including cell-killing assay, were performed following the published methods (28, 30). Further details of the materials and methods used in this study are described in *SI Appendix*.

## ACKNOWLEDGMENTS

We thank NYU Langone Health’s Cryo–Electron Microscopy Laboratory (RRID: SCR_019202) for cryo-EM work and Dr. William Rice and Dr. Bing Wang for their technical assistance. We also thank Dr. Brian D. Dynlacht for access to the Incucyte live-cell analysis system. This work was supported by National Institutes of Health grants R21CA246457 (T.H.), R01CA251669 (C.Y.P. and S.K), R01NS102665 (D.G.P.), R01NS124920 (D.G.P.), R21NS126806 (D.G.P.) and

P30CA016087 (Cancer Center Support Grant).

## DATA DEPOSITION

The maps and structure data have been deposited in the Electron Microscopy Database and PDB (EMD-48537 and 9MQR).

## AUTHOR CONTRIBUTION

T.H., M.W., A.D.C., S.G., D.G.P., C.Y.P and S.K. designed research. T.H., M.W., A.D.C., S.G., M.F., I.B., N.R., I.B., H.D., C.G., N.R.B. and A.K. performed research. All authors analyzed data. T.H., M.W., S.G., D.G.P., C.Y.P and S.K. wrote a paper with input from other authors.

